# Aberrant Sensory Encoding in Patients with Autism

**DOI:** 10.1101/2020.03.04.976191

**Authors:** Jean-Paul Noel, Ling-Qi Zhang, Alan A. Stocker, Dora E. Angelaki

## Abstract

Perceptual anomalies in patients with Autism Spectrum Disorder (ASD) have been attributed to irregularities in the Bayesian interpretation (i.e., decoding) of sensory information. Here we show that how sensory information is encoded and adapts to changing stimulus statistics also characteristically differs between healthy and ASD groups. In a visual estimation task, we extracted the accuracy of sensory encoding directly from psychophysical data, bypassing the decoding stage by using information theoretic measures. Initially, sensory representations in both groups reflected the statistics of visual orientations in natural scenes, but encoding capacity was overall lower in the ASD group. Exposure to an artificial statistical distribution of visual orientations altered the sensory representations of the control group toward the novel experimental statistics, while also increasing their total encoding resources. Neither total encoding resources nor their allocation changed significantly in the ASD group. Most interestingly, across both groups the adaptive re-allocation of encoding resources was correlated with subjects’ initial encoding capacity. These findings suggest that neural encoding resources are limited in ASD, and this limitation may explain their reduced perceptual flexibility.

## Introduction

Recent theories attempting to provide a normative account for the complex phenotype of Autism Spectrum Disorder (ASD) have used the language of statistical inference, proposing that prior knowledge about the world is under-emphasized relative to incoming sensory information in patients with ASD. The primary source of this imbalance is debated: some authors argue for attenuated priors (Karaminis et al., 2016; Pellicano and Burr, 2012a; 2012b; Powell et al., 2016), while others argue for aberrant sensory precision or prediction error (Karvelis et al., 2018; Lawson et al., 2014; Palmer et al., 2017; Brock, 2012; Van de Cruys et al., 2013). In common, however, these studies are disproportionally emphasizing the interpretation of incoming sensory information (i.e., decoding) while ignoring the possibility that deficits in the neural representation of sensory information (i.e., encoding) may also be responsible for the observed differences in perceptual behavior between ASD and healthy populations.

Here we focused on characterizing sensory encoding in ASD while purposefully remaining agnostic about the decoding process. The goals were to isolate the contribution of changes in encoding from potential effects attributed to deficits in decoding. To examine the capacity and flexibility of sensory encoding in ASD we asked participants to perform a visual orientation estimation task, first without feedback, and then in the presence of trial-by-trial feedback. We took advantage of the well-known “oblique effect” (Campbell et al., 1996; Appelle, 1972) where humans demonstrate greater sensitivity and repulsive (i.e., away from) biases in perceiving orientation and motion at cardinal compared to oblique orientations (Westheimer & Beard, 1998; Dakin et al., 2005). These effects can be explained by postulating that the visual system efficiently encodes sensory information (Wei & Stocker, 2015, 2017) - i.e., by an encoding stage that is matched to the natural environmental statistics where vertical and horizontal edges are most common (Coppola et al., 1998, Girshick et al., 2011). In other words, the distribution of orientations in natural scenes imposes a sensory representation that allocates less sensory resources for low probability stimulus values, resulting in a less accurate neural representation of these stimuli.

We analyzed the psychophysical estimation data using an information theoretic approach, the Cramer-Rao bound, that leverages the lawful relation between estimation bias, variance, and encoding accuracy (Casella & Berger, 2002; Wei & Stocker, 2017). The approach allows us to determine the accuracy of sensory representation while making only the minimal assumption that the efficiency of the decoder is independent of stimulus orientation. From measurements of perceptual biases and variance we estimated a lower bound of Fisher Information (FI), a measure of encoding accuracy. Subsequently, based on the efficient coding hypothesis (Wei & Stocker, 2015), we can further postulate that encoding accuracy reflects subjects’ expectations of the stimulus distribution (i.e., their prior). We express these expectations as a combination of the distribution of orientations in natural scenes (Coppola et al., 1998, Girshick et al., 2011) and the uniform distribution of orientations imposed by the experiment. The initial trials in the absence of feedback allowed us to compare overall encoding capacity and its allocation as a function of orientation between ASD and control groups. The second set of trials allowed us to examine how both properties changed in the presence of feedback. Our results show encoding capacity limitations and a lack of encoding flexibility in subjects with ASD. These findings suggest that perception in patients with ASD may not only anomalous in the way sensory information is interpreted, but crucially also in the accuracy and flexibility of how sensory information is encoded in the first place.

## Results

### Heightened variability and reduced learning in Autism

Groups of ASD (n = 17) and control (n = 25) individuals matched a briefly (120ms) presented visual target of random orientation. Target orientations were drawn from a uniform distribution. Initially no feedback was presented (woFB block), but in the second and third block of trials (wFB1 and wFB2; 200 trials/block) feedback was presented by overlaying the participant’s response and the target (**Fig 1A**, see *Methods* for details). As shown with an example control (**Fig. 1B**) and ASD (**Fig. 1C**) individual, orientation perception was biased away from cardinal orientations in both groups – namely, both groups demonstrated an “oblique effect” (Campbell et al., 1996; Appelle, 1972). With feedback, the bias seemingly dissipated in the control subject but not the ASD individual (**Fig. 1B, C** scatter plot are individual responses, curve is a running average within a sliding 15° Gaussian kernel, and vertical dashed lines are orientations at 45°, 90°, and 135°. See **Fig. S1** for similar plots for all individual subjects).

**Figure 1.**
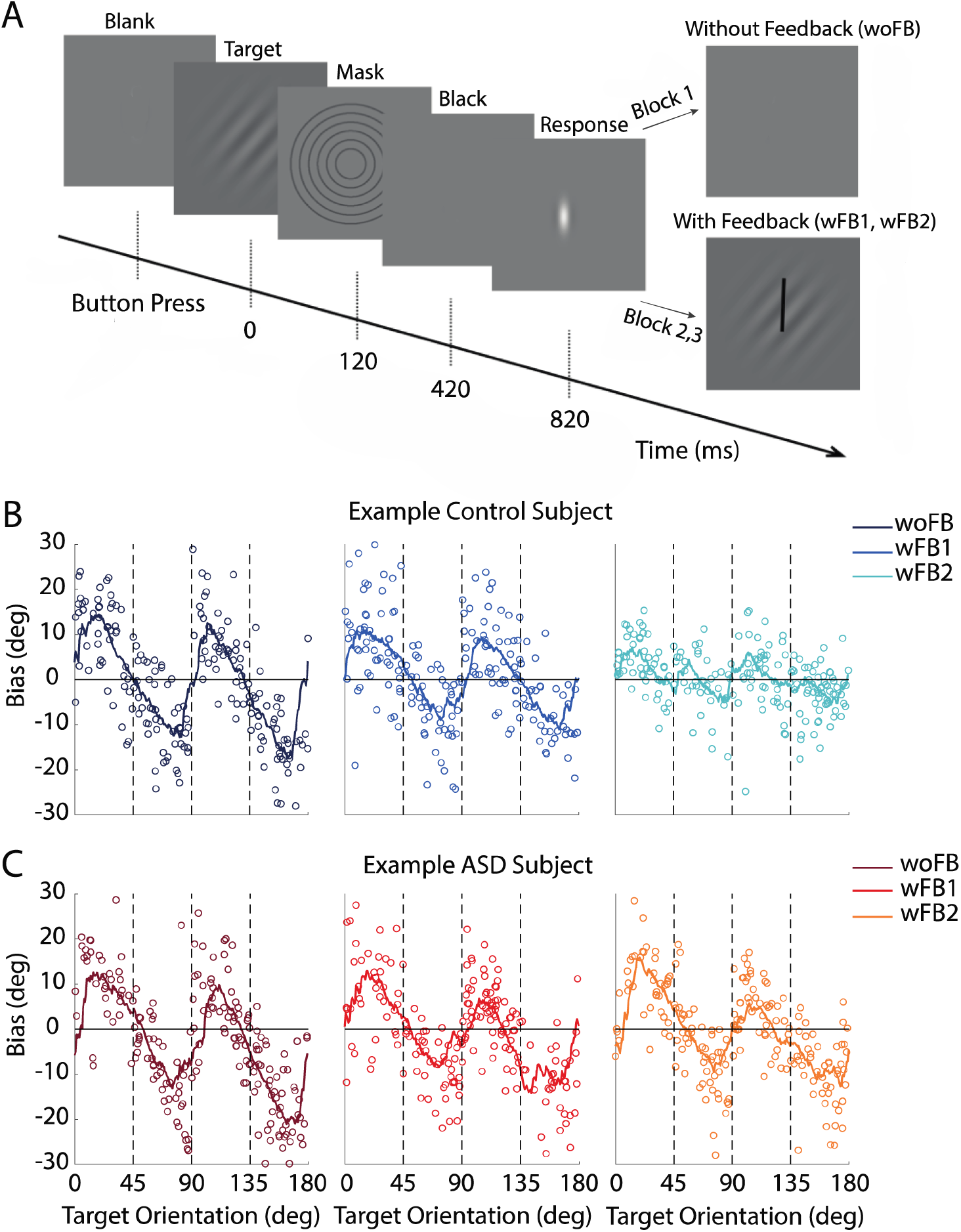
Experimental Protocol and Individual Subject Performance. **A)** A target orientation (Gabor) is briefly presented and participants report their percept by orienting a line indicator (white). No feedback is given on the first block of trials, but is in subsequent blocks by overlaying the target orientation and the participant’s response. **B)** Target orientations (x-axis) are drawn from a uniform distribution (individual dots are single trials). Y-axis indicates the bias for an example control subject (response subtracted from target), and lines are the running average within a sliding Gaussian kernel of 15°. Different columns (and the color gradient) respectively show performance on the block without feedback (woFB; leftmost), on the first block with feedback (wFB1; center), and the second block with feedback (wFB2; rightmost). **C)** follows the format in **B)** while depicting the performance of an example ASD subject.

These basic observations are also evident in group averages of control and ASD individuals (**Fig. 2**). When presenting targets between 0° (horizontal) and 45°, the bias was on average positive (e.g., average bias for control group: 6.0° ± 0.26° (S.E.), p < 10^−3^), suggesting that horizontal gratings were perceived closer to the oblique 45°. Contrary, when presenting targets between 45° and 90° (vertical), the bias was negative (e.g., average bias for control group: -5.14° ± 0.26°, p < 10^−3^), suggesting that vertical gratings were also perceived closer to the oblique 45° (**Fig. 2A, B**). Before feedback, there was no statistically significant difference in the overall magnitude (i.e., absolute value) of bias between control and ASD groups (control: 5.63° ± 0.12°; ASD: 5.99° ± 0.22°, Δ = 0.35° ± 0.25°, p = 0.077). On the other hand, when provided with feedback, orientation perception bias was reduced in the control group, but less so in the ASD group (reduction in average magnitude of bias between the block without feedback (woFB) and the second set of trials with feedback (wFB2), control: Δ = 2.38° ± 0.17°, p < 10^−3^; ASD: Δ =1.04° ± 0.31°, p < 10^−3^; ΔControl - ΔASD = 1.34° ± 0.35°, p < 10^−3^; **Fig. 2A, B**, feedback conditions shown by a color gradient).

**Figure 2.**
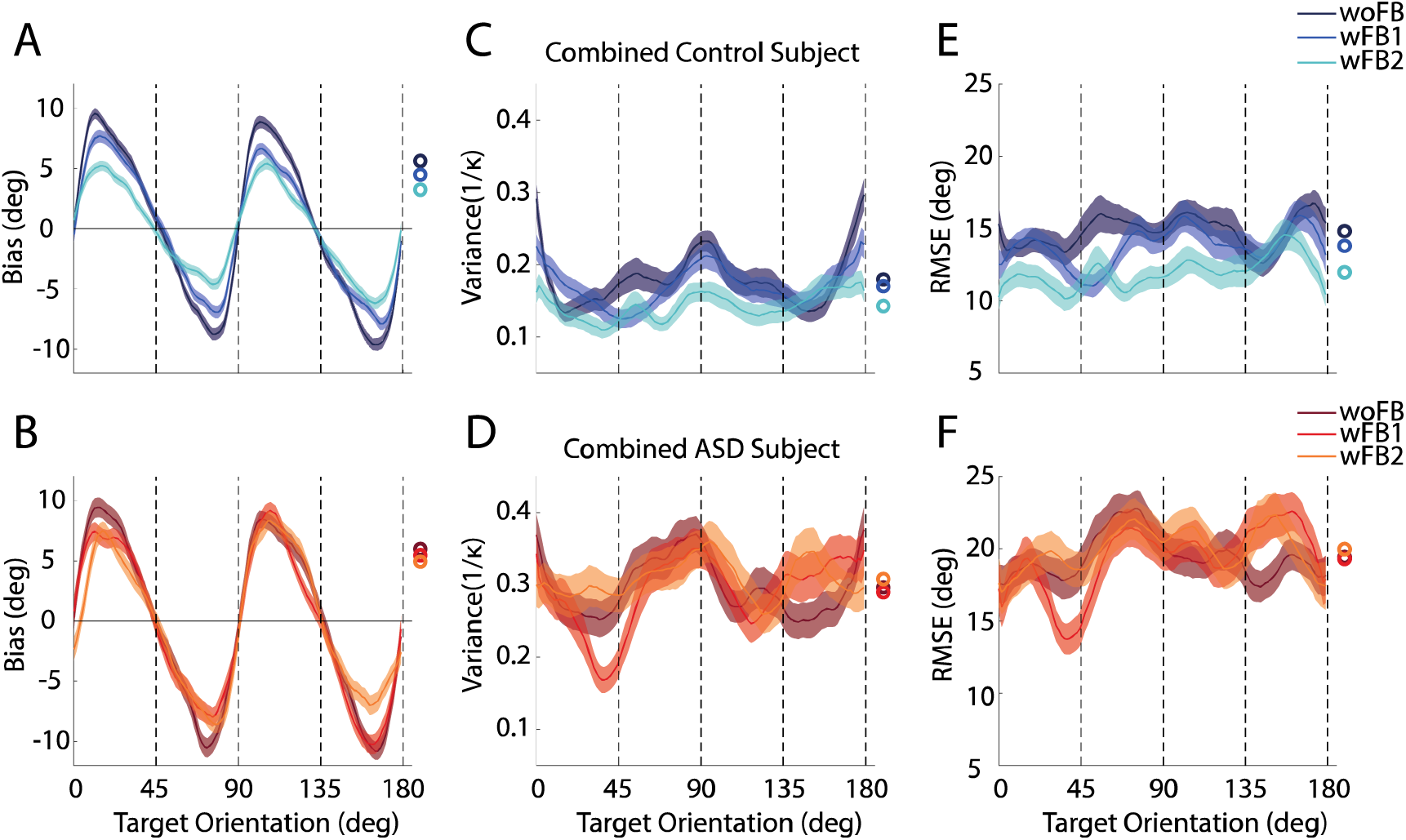
Orientation Perception in Combined Control and ASD Subject. Bias (y-axis) as a function of target orientation (x-axis, cardinal and oblique orientation indicated by dashed lines) and feedback block in control **(A**) and ASD **(B)** subjects. Variance (1/kappa, y-axis) as a function of target orientation and feedback block in control **(C)** and ASD **(D)** subjects. Root-mean-square error (RMSE, y-axis) as a function of target orientation and feedback block in control **(E)** and ASD **(F)** subjects. Error bars are ± SEM across 5,000 bootstrap runs.

Regarding the variability of orientation perception, we found that at baseline (i.e., before feedback), the ASD group had larger variance in their estimates than the control group (control: 0.180 ± 0.006; ASD: 0.297 ± 0.010; Δ = 0.117 ± 0.11, p < 10^−3^; **Fig. 2C, D**). This is in line with a growing literature suggesting heightened sensitivity to noise in ASD (e.g., Dinstein et al., 2012; Haigh et al., 2014; Zaidel et al., 2015; Noel et al., 2019). Additionally, while variance was seemingly further reduced with feedback in the control group, especially in the latter feedback block, this was not evident in the ASD cohort (reduction in overall variance between woFB and wFB2, control: Δ = 0.037 ± 0.008, p < 10^−3^; ASD: Δ = -0.012 ± 0.015, p = 0.414).

As expected, based on these differences in bias and variability, the control group overall had better performance as measured by root-mean-square error (RMSE). This was true in the initial block of the experiment (control: 14.82° ± 0.35°, p < 10^−3^; ASD: 19.42° ± 0.47°, p < 10^−3^), and was exacerbated with feedback (**Fig. 2E, F**). Namely, performance of the control but not the ASD group increased with feedback, (reduction in overall RMSE between woFB and wFB2, control: Δ = 2.83° ± 0.48°, p < 10^−3^, ASD: Δ = -0.55° ± 0.71°, p = 0.78).

### Extracting accuracy of sensory encoding from psychophysical estimation data

The observed differences in bias and variance between ASD and control individuals (**Figs. 1, 2**) could originate from differences in the encoding process (i.e., how a target is sampled in noisy neural representations) or as a growing literature emphasizes, in how the sensory representations are interpreted in light of prior knowledge and expectations (e.g., Pellicano & Burr, 2012) (**Fig. 3A**, encoding step vs. decoding step). To disambiguate between these possibilities some (e.g., Karvelis et al., 2018) have recently jointly fit likelihood functions and priors to psychophysical data from individuals with ASD. This exercise allows estimating the shape of a likelihood function and one may equate the latter with sensory encoding. However, this approach is problematic as it 1) assumes an intact inference process in ASD (Palmer et al., 2017), and 2) ignores more nuanced and interdependent relations between Bayesian priors, sensory likelihoods, and posteriors beyond that specified by Bayes’ Rule (posterior = likelihood x prior). Namely, as suggested by efficient coding, the prior may directly constrain the shape of likelihood distributions (e.g., Wei & Stocker, 2015), or as suggested by hierarchical Bayesian models (e.g., Mathys et al., 2014), posteriors at one level may be the likelihood functions at the next. Further, during the feedback condition subjects may develop explicit and idiosyncratic strategies to reduce error in their estimate that cannot be fully captured by a restricted family of estimators, such as Bayesian decoders.

**Figure 3.**
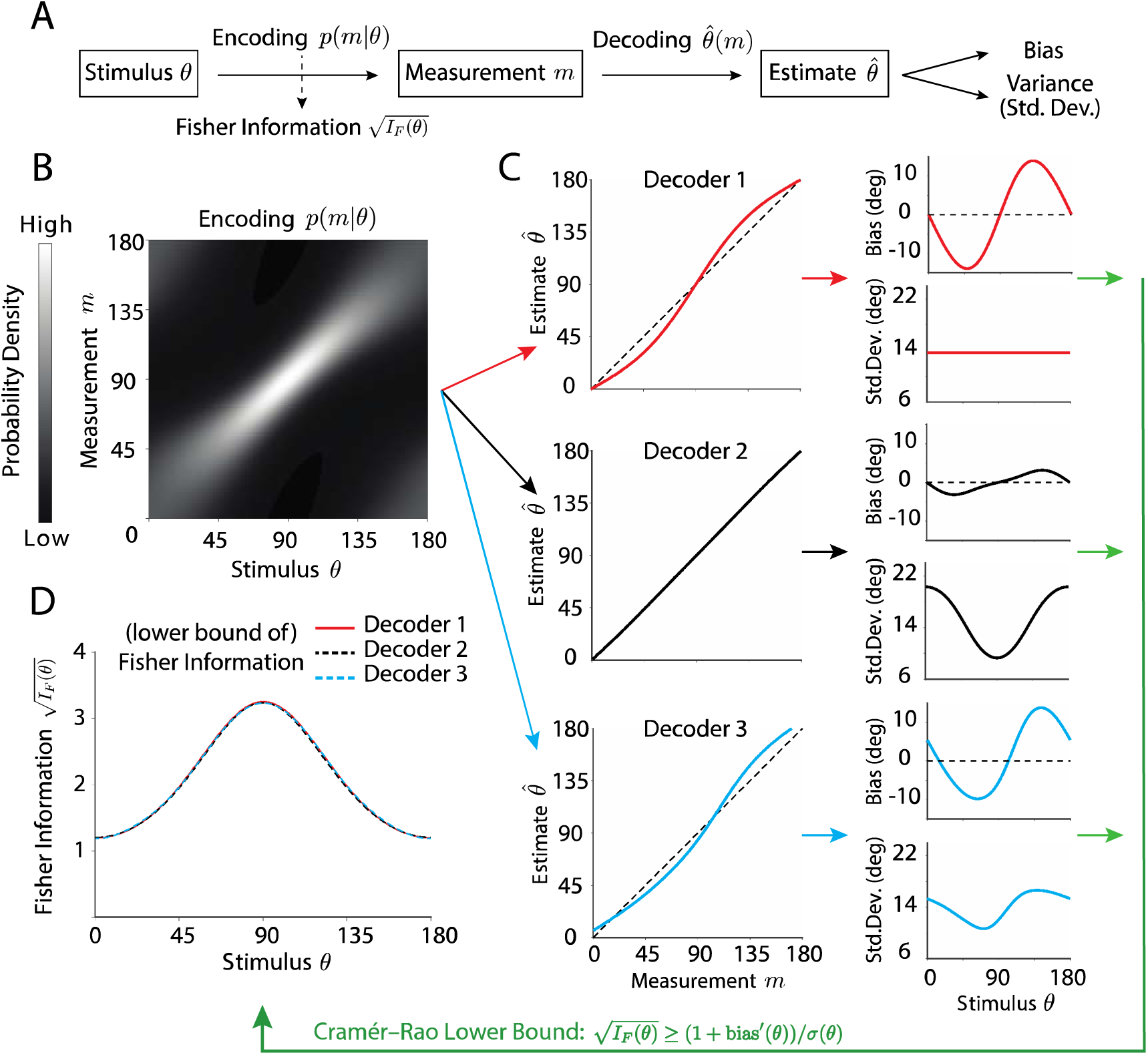
Simulation Demonstrating that Fisher Information is a Measure of Sensory Encoding Accuracy that is Independent of the Decoding Process. **A)** Perception can be described by a process whereby an external stimulus θ, is first encoded by a noisy measurement, p(m|θ), which is then decoded into an estimate 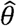. The estimate can be characterized by a bias and variance in perception. **B)** To examine sensory encoding independently of the decoding process, we use the lawful relation between (the derivative of) bias and variance to estimate Fisher Information (FI). The latter being a measure of encoding accuracy, as we illustrate by a simulation. We assume an encoding process, p(m|θ), with a peak FI at 90°. **C)** As an example, we construct three arbitrary decoders (red, black, and blue) yielding different biases and variances. **D)** Applying the Cramer-Rao Inequality we estimate the lower bound of FI, which appropriately suggests that all three patterns of bias and variance possess the same pattern of FI.

To overcome these issues, our goal here was to return to more basic concepts and to examine neural encoding within a normative framework that is independent from the decoding process. This may inform whether purported deficits in the decoding process may be inherited from anomalous encoding. For this, we leverage a lawful relationship between the bias and variance of an estimator, the Cramer-Rao lower bound (Casella & Berger, 2002; Wei & Stocker, 2017), to estimate Fisher Information (FI). This latter measure describes how much information subjects’ internal representation carries about the physical stimulus presented, and provides a lower bound for the discrimination threshold of an observer based on that representation (Seung & Sompolinsky, 1993; Series et al., 2009). To illustrate the independence of this approach from the specific decoders, we present a simulation. We assume an encoding process p(*m*|*θ*) with a peaked (von mises) FI centered around 90° (**Fig. 3B**). Then, we built three arbitrarily chosen decoders 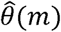 which given a sensory measurement *m*, generate an estimate 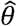 (**Fig. 3C**). These decoders each produces a distinct pattern of bias and variance in their estimate, albeit based on the same encoding process (**Fig. 3C**). The Cramer-Rao lower bound (**Eq. 1**) states that, the bias and variance pattern produced by any decoder based on the same encoding process, has to satisfy the relationship (see *Methods*):

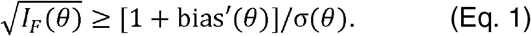

Thus, upon observing the bias and variance, we can estimate a (lower bound) of FI in the encoding process (**Fig. 3D**). As example, despite the distinct behavior of the three decoders in this stimulation, we can extract the exact same actual pattern of FI in the encoding process. The only assumption here is that the decoder is efficient (i.e., it achieves the bound), which we verified as a valid assumption for a large range of decoders (also see *Discussion*). In turn, we use the Cramer-Rao Inequality and FI to examine encoding capacity and flexibility in ASD and control individuals.

### Reduced capacity and aberrant allocation of encoding resources in Autism

Two aspects of FI are important for characterizing the sensory encoding of our subjects: 1) the overall scale, which determines the total amount of encoding resources, and 2) the shape or pattern of FI, which determines how resources are allocated among different orientations.

Regarding the shape of FI as a function of target orientations, we found that in both groups FI peaked at cardinal orientations, those that are most common in the natural environment (**Fig. 4A**). The overall (normalized) pattern of FI matches the previously measured distribution of orientations in natural images (Dong & Atick, 1995; Roth & Black, 2005; Coppola et al., 1998, Girshick et al., 2011). On the other hand, both the total amount of FI and how this measure changed over the course of the experiment differed for ASD and control groups. Already during the first block of trials (woFB), total FI was significantly lower in ASD than controls (woFB, control: 14.67 ± 0.24, ASD: 11.28 ± 0.19). Further, total FI increased from woFB to wFB2 for the control group (Δ = 1.96 ± 0.38, p < 10^−3^), but did not change in the ASD group (Δ = -0.060 ± 0.27, p = 0.83, **Fig. 4B**).

**Figure 4.**
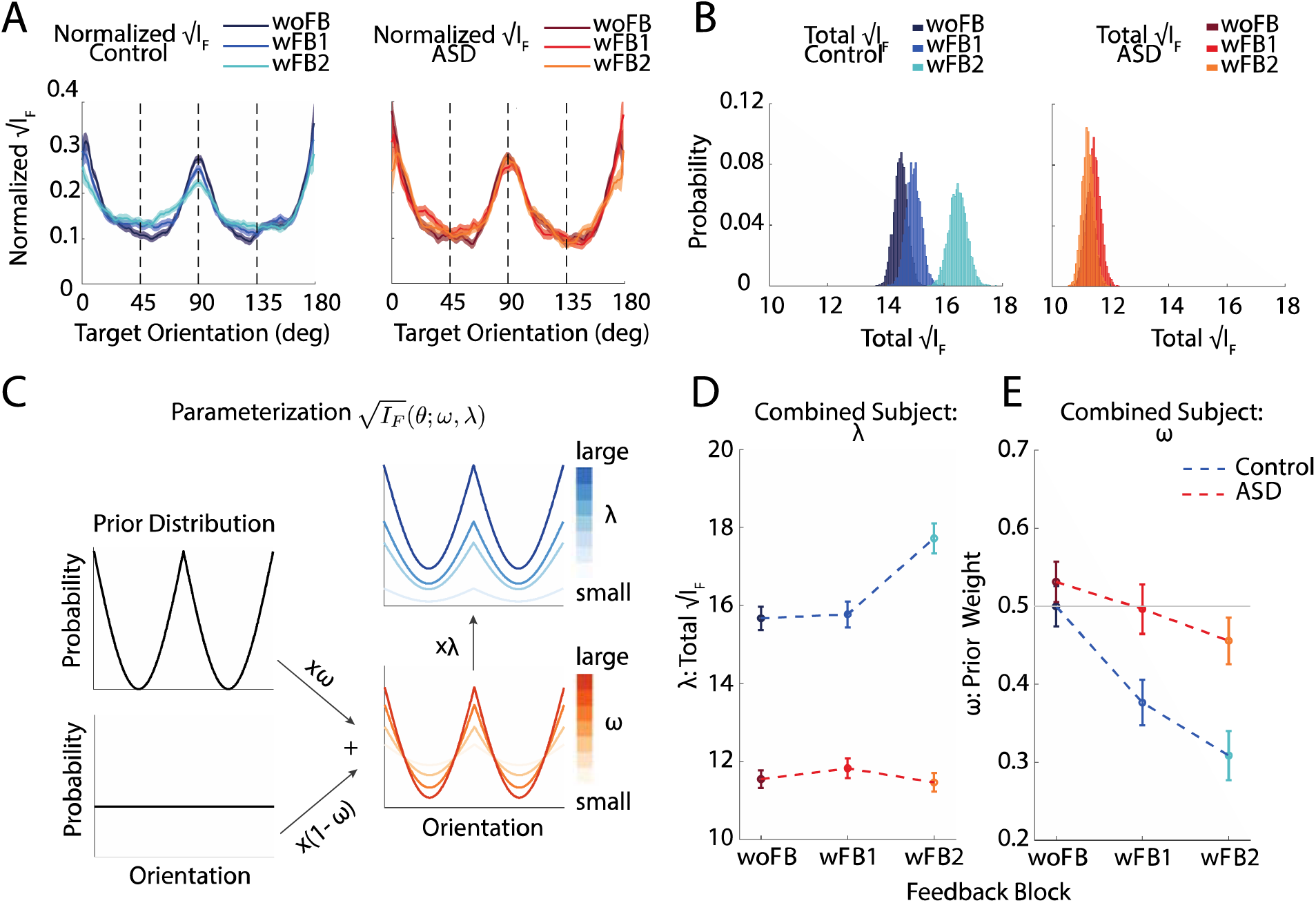
Quantification and Parametrization of FI in Control and ASD individuals. **A)** Fisher Information (FI) peaked at cardinal orientations for both control (blue) and ASD (red) individuals. **B)** The total amount of FI was larger in control than ASD at the outset, and increased over the course of the experiment in control (blue color gradient) but not ASD (red color gradient) individuals. **C)** The FI pattern as a function of orientation was quantified by two parameters, ω, which mixes a natural orientation prior with a uniform distribution as the normalized (square-root of) FI, and λ, scaling total FI. **D)** λ and **E)** ω as a function of group (blue = control, red = ASD) and block. Note a ω value of 0.5 (grey line) roughly corresponds to what has been previously reported as the prior distribution of orientations in natural environment. Error bars are ± SEM across 5,000 bootstrap runs.

To more fully quantify the impact of feedback on the shape and amount of FI, and more importantly, to relate our information theoretic measure (FI) to previous Bayesian accounts (e.g.,Pellicano & Burr, 2012; Lawson et al., 2017; Karvelis et al., 2018), we parametrized the normalized square root of FI as a weighted sum of the distribution matching the known statistics for orientations in the environment (Girshick et al., 2011) and the experimental uniform distribution (with weights *ω* and 1-*ω*, respectively; **Fig. 4C**, Eq. 3-6 in *Methods*). That is, under the framework of efficient coding the square root of FI is proportional to the stimulus distribution, the prior (Eq. 2, also see Wei & Stocker, 2015). Efficient coding dictates that sensory system ought to allocate encoding resources to match the natural statistics of the environment and under this theoretical framework, a flattened FI (**Fig. 4A**, controls) is expected if subjects begin to adapt their natural prior in order to incorporate the uniform prior imposed by the experiment (given that random orientations presented in the current experiment were drawn from a uniform distribution). The smaller *ω*, the closer the participants’ prior to the statistics imposed by the experiment. Further, the overall scale of the square root of FI (i.e., the amount of total FI) is determined by another parameter, λ (**Fig. 4C**). We then fitted the bias predicted by the parameterized FI to the measured bias of the combined subject, given the measured variance (see *Methods* and **Fig. S1** for individual participant biases and fits). That is, differently from previous computational work within the study of ASD, the prior (i.e., square root of FI) here is completely constrained by the participants’ responses (i.e., variance and bias, according to the Cramer-Rao Inequality).

Examination of the fitted λ values confirmed that the ASD group had a lower baseline total (square-root of) FI than the control group (control λ vs. ASD λ at woFB; Δ = 4.121 ± 0.385, p < 10^−3^; **Fig. 4D**). Over the course of the experiment, λ further increased for the control group (woFB vs. wFB2 in control: Δ = 2.039 ± 0.502, p < 10^− 3^; **Fig. 4D** but this was not the case for the ASD group (Δ = -0.088 ± 0.325, p = 0.788, **Fig. 4D**). These results suggest an aberrant encoding process in the ASD group, showing a lower overall encoding capacity that does not improve over the course of repeated feedback.

Notably, both groups started with a similar orientation-dependent pattern of normalized FI, with no significant difference in the *ω* parameter prior to feedback (Δ = 0.032 ± 0.037, p = 0.391, Fig. **4E**). When provided with feedback (woFB vs. wFB2), the *ω* parameter decreased, thus the overall FI pattern became flattened, for the control group (change in *ω*: Δ = 0.191 ± 0.040, p < 10^−3^, **Fig. 4E**, blue), but much less so for the ASD group (change in *ω*: Δ = 0.076 ± 0.039, p = 0.027, contrast between control and ASD: Δ = 0.114 ± 0.057, p = 0.020). This change in the *ω* parameter under the efficient coding framework can be viewed as control subjects altering their prior of orientations by incorporating the uniform prior imposed by the experiment, compared to a less sensitive ASD group. That is, with feedback not only did the total amount of encoding resources increased in control but not ASD individuals, but the allocation of these resources also adapted to the experimental requirements in the control and much less in the ASD group.

### Reduced initial Encoding Capacity Is Correlated with flexibility in Prior

Recent theoretical studies (Mlynarski & Hermundstad, 2018, 2019) highlight an inherent relation between encoding resources and the degree to which one can adapt to changing environments. In other words, if in line with efficient coding (Barlow, 1961; Wei & Stocker, 2015) resources are allocated primarily to represent statistically likely events, then fewer can be devoted to distinguishing between statistically unlikely alternatives – potentially limiting one’s knowledge that the environment has changed or how (Mlynarski & Hermundstad, 2018, 2019). To examine this hypothesis, we repeated the analysis above for each individual subject and correlated participants’ encoding capacity (λ) before feedback with the shape of their prior after feedback (*ω* at wFB2). In line with the “adaptive coding” hypothesis stated above (Mlynarski & Hermundstad, 2018, 2019), we found a strong correlation between these variables (r = 0.371, p < 0.001, **Fig. 5A**). Thus, individuals with ASD may have inflexible priors because of their limited encoding resources (see *Discussion*).

**Figure 5.**
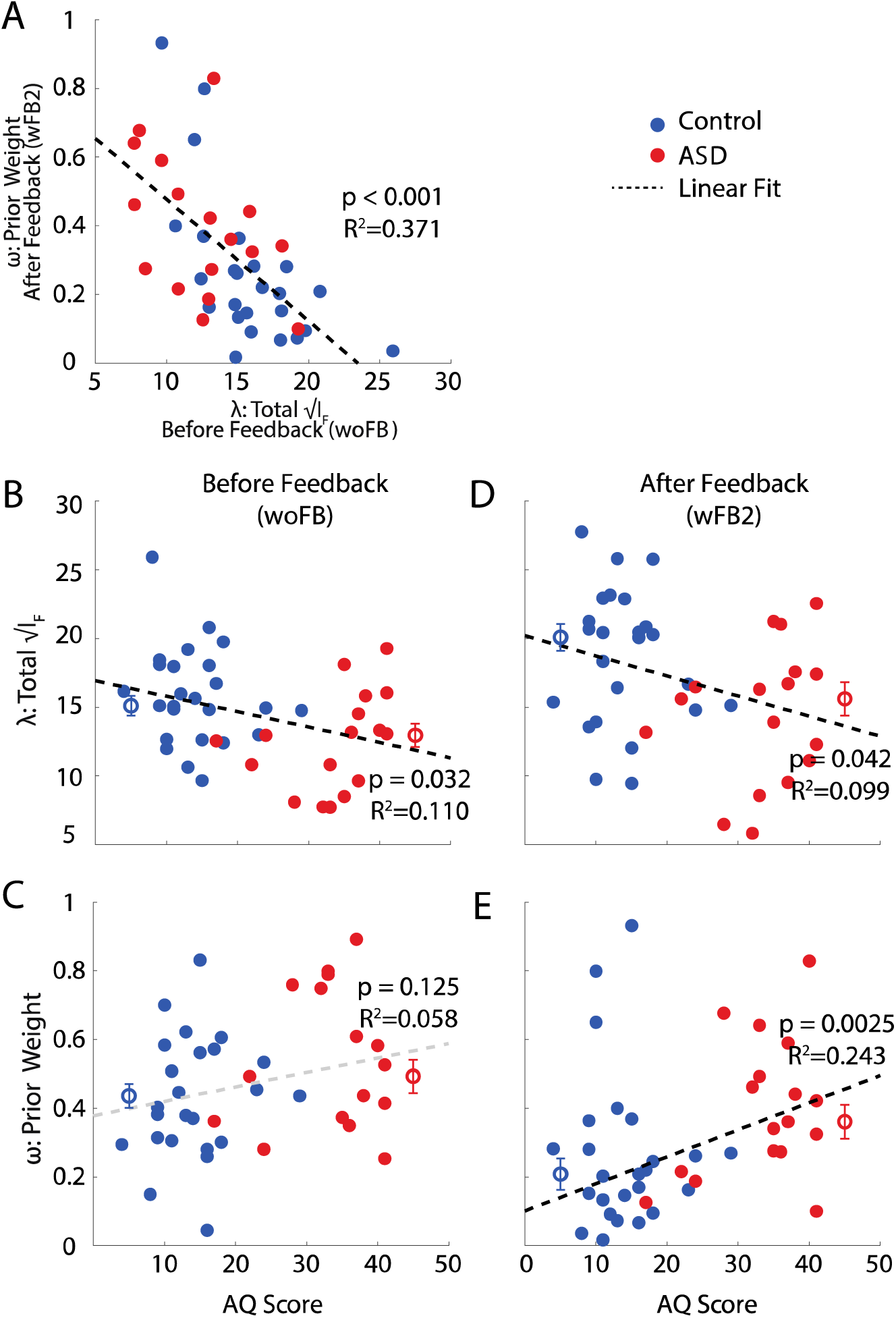
Correlation between Sensory Encoding Capacity, Flexibility in Encoding Resources Allocation, and Measures of Autism Traits. **A)** The flatness of FI after feedback (y-axis, indicating incorporation of stimulus statistics from the experiment) correlates with total encoding resources before feedback (x-axis). The total amount of encoding resources (y-axis) both before **(B)** and after **(C)** feedback correlates with Autism Quotient Scores. The shape of the prior before feedback **(D)** does not correlate with Autism trait severity, but does after feedback **(E).** Controls are depicted in blue, and ASD subjects are shown in red (individual dots are single participants, dots with errors bars are means across individuals ± SEM). Dark dashed lines are regression slopes that are significant, while the gray dashed line is not.

Further supporting a potential driving role of encoding capacity in the reduced flexibility of priors in ASD, we also found that the total FI (λ) before feedback, but not the shape of the prior distribution (*ω*), correlated with ASD symptomatology, as indexed by the Autism Quotient (AQ, Baron-Cohen et al., 2001; λ and AQ: r = 0.110, p = 0.032; *ω* and AQ: r = 0.058, and p = 0.125, **Fig. 5B, C**). On the other hand, after feedback (wFB2) both λ (r = 0.099, and p = 0.042) and most importantly, *ω* (r = 0.243, and p = 0.0025, **Fig. 5D, E**) – potentially driven by λ before feedback - showed a significant correlation with the AQ. Similar correlations existed as well with the Social Communication Questionnaire (Rutter et al., 2003; see **Fig. S2**).

## Discussion

By 1) tracking perceptual estimates over changes in stimulus statistics, 2) directly extracting Fisher Information (FI) from psychophysical data, and 3) parameterizing the shape of FI as a function of the stimulus and natural distributions, we have estimated the total encoding capacity and the allocation thereof, a proxy for the prior within the efficient coding framework, in individuals with ASD. We show that neural encoding resources are limited in ASD and suggest that this limitation may explain their aberrant perceptual flexibility (e.g., Lawson et al., 2017; Leider et al., 2019).

We found that the estimated natural prior for visual orientations is initially equal across autistic and neurotypical populations, yielding similar orientation-matching biases. Initially, the main difference between groups was the lower encoding capacity in ASD, reflected behaviorally as a higher within-subject variability (Zaidel et al., 2015; Noel et al., 2019). Using a normative framework, we were able to estimate that, at least within the current orientation estimation task, the total amount of encoding resources within the ASD group was about 75% that of their neurotypical counterparts. Further, the total amount of encoding resources correlated significantly with autistic symptomatology. This is important since recent simulations (Mlynarski & Hermundstad, 2018, 2019) highlight an inherent relation between encoding resources and the degree to which one can adapt to changing environments. Namely, in line with efficient coding (Barlow, 1961; Wei & Stocker, 2015) where resources are allocated primarily to represent statistically likely events, fewer resources may be devoted to distinguishing between statistically unlikely alternatives – thus, potentially limiting one’s knowledge that the environment has changed or how.

Indeed, we found that the degree to which the encoding prior changed after feedback was correlated with initial encoding capacity: participants with greater initial encoding capacity had a prior that is closer to a uniform (experimental) distribution at the end of the experiment. Further, we also found that only the control group adapted their allocation of encoding resources from matching the statistics of the natural environment to matching those of the experiment. This was not observed in the ASD group. Further, the change in the pattern of allocating encoding resources correlated with autistic symptomatology. The only measure not correlating with ASD symptomatology was the prior before feedback, arguably the only parameter not dependent on encoding resources. Together, these findings argue for the importance of sensory encoding limitations in autistic phenotypes. Given that the prior was initially similar across groups, but FI was not, we argue that reduced encoding capacity leads to the loss of flexibility and adaptability in the ASD group. Thus, our findings suggest that the reason priors are inflexible in ASD may be because of a limited pool of encoding resources, and thus the inability to represent statistically unlikely events.

In recent years, several ASD studies have employed a Bayesian decoder framework and have argued for weaker and/or inflexible priors in the condition (e.g., Lawson et al., 2017; Lieder et al., 2019; Powell et al., 2016; Palmer et al., 2017). Our results do not contradict these conclusions, they simply emphasize that a computational understanding of the ASD phenotype cannot be limited to Bayes’ Rule (Posterior = Prior * Likelihood). In fact, the observed reduction of FI in ASD may be in agreement with the hypo-prior conjecture: expectations – particularly if they are accurate priors – may effectively implement a low-pass filter (e.g., moving average) reducing variance and uncertainty. In this perspective the panoply of studies demonstrating heightened variability (Distein et al., 2012; Haigh et al., 2014; Bonneh et al., 2011; Milne, 2011; Noel et al., 2019) and heightened sensitivity to sensory noise (Zaidel et al., 2015) in ASD are all in line with either the hypo-prior account or a reduction in encoding resources.

However, Bayesian decoding analysis neglect that, before interpreting sensory representations by decoders, environmental signals must be captured by noisy sensors at the sensory periphery and encoded by stochastic neurons. To account for encoding independently from decoding, here we have examined for the first time sensory representations in ASD within a principled framework. The computational choice is supported by the fact that the estimated FI matches the known distribution of orientations in the environment, and our analysis has the advantage that it only makes the assumption that the decoder is efficient (i.e., the decoder is at the Cramer-Rao lower bound). A wide range of decoders may apply, and we need not assume a particular decoder (see simulations in **Fig. 3**). This is important, emphasizing that previous studies concluding a particular deficit in the priors or likelihood functions of a decoder have assumed appropriate Bayesian decoding in ASD, which has never been explicitly tested.

Nonetheless, one may wonder whether the reported effects may be due to differential efficiency of decoders in the control and ASD population. In a set of additional simulations (**Fig. S3**), we show that decoding inefficiency that is independent of the stimulus (e.g., homogeneous noise added to the decoder) will both decrease the total amount of FI and flatten the pattern of FI. This is incompatible with our results in two aspects. First, the control and ASD group differ only in the total FI (i.e., the λ parameter), but not the pattern of FI (i.e., the *ω* parameter) before receiving feedback. Second, learning through feedback increases total FI while flattening the pattern of FI. If the increase in total FI were to be explained by an increase in the efficiency in the decoder, the pattern of FI ought to be sharpened, which is the exact opposite of what we observed in the data.

There is a wide range of potential neural mechanism by which the amount and allocation of encoding resources may change relatively rapidly. A first possibility is a gain modulation of the neural response. Indeed, for an efficient coding framework where each cell transmits an equal portion of the stimulus probability mass, an increase in the overall firing rate corresponds to a direct increase in population FI (Ganguli & Simoncelli, 2014). Gain modulation could be implemented by alterations in neuromodulation, and fittingly Lawson and colleagues (2017) recently demonstrated abnormal noradrenergic responsivity in ASD. Previous studies of sensory adaption (e.g., Clifford et al., 2007) have also suggested that gain changes specific to a subpopulation of neurons are able to alter the allocation of coding resources. A second potential mechanism may be changes in the structure of correlated noise in the neural population. For example, a decrease in interneuronal correlations generally reduces the impact of noise on stimulus representation, thus leading to a higher population FI (Cohen & Maunsell, 2009). Along this line, Coen-Gagli & Solomon (2019) have recently suggested that divisive normalization is a critical player in neural variability, and Rosenberg and colleagues (2015) have accounted for a wide array of perceptual deficits in ASD by postulating a deficit in divisive normalization.

In conclusion, recent theories within computational psychiatry have proposed that a range of psychopathological disorders, most notably Autism (Pellicano & Burr, 2012; Van der Cruys et al., 2014; Lawson et al., 2014) but also Schizophrenia (Fletcher & Frith, 2009; Corlett et al., 2009; Adams et al., 2013), are underpinned by deficits in performing Bayesian inference: the ability to correctly integrate sensory information with prior knowledge. These psychiatric-focused Bayesian theories, however, have so far largely ignored the possibility that the encoding of sensory information itself could be affected (but see Karaminis et al., 2016; Karvelis et al., 2018). They generally assumed that the generation of likelihood functions (encoding) and its combination with a prior (decoding) are independent processes. However, such an assumption runs counter the efficient coding framework (Barlow, 1961) and the normative foundation that the nervous system is optimized to represent those stimuli it most often encounters. Having taken this property into account, the present analyses suggest that in ASD a reduced pool of encoding resources leads to increased variability and the inability to accurately represent statistically unlikely events. This may explain the observed inflexibility in the re-allocation of sensory resources. More importantly, our results suggest that the encoding stage itself appears aberrant in ASD. Thus, future studies must carefully characterize other aspects of normative computation, like efficient coding, before concluding that it is the (Bayesian) decoding phase that has gone awry in ASD.

## Online Methods

### Participants

A total of forty-two subjects completed an orientation-matching task. Seventeen were individuals diagnosed as within the Autism Spectrum Disorder (ASD; N = 17, mean ± s.d.; age = 15.3 ± 2.6 years; AQ = 33.5 ± 7.0; SCQ = 16.8 ± 4.4) by expert clinicians according to the Autism Diagnostic Observation Schedule (ADOS; Lord et al., 2000). The rest were neurotypical individuals (Control; N = 25, mean ± s.d.; age = 14.8 ± 2.1 years; AQ = 14.0 ± 5.5; SCQ = 5.6 ± 3.2). Participants had normal or corrected-to-normal vision, and no history of musculoskeletal or neurological disorders. Before partaking in the study, all participants completed the Autism Spectrum Quotient (AQ; Baron-Cohen et al. 2001) and the Social Communication Questionnaire (SCQ; Rutter et al, 2013). The Institutional Review Board at Baylor College and Medicine approved this study, and all participants gave their written informed consent and/or assent.

### Materials and Procedures

Participants were comfortably seated facing a gamma-corrected CRT monitor (Micron Technology, Boise, ID; 43 × 35 cm) at a distance of 57 cm. Subjects self-initiated a trial by button press, upon which a Gabor (120ms presentation, 0.4 cycles/degree, 5 cm radius, Gaussian envelope) was presented centrally on a gray background. The orientation of this target Gabor was random (uniform distribution, 0 – 180 degrees). Immediately following the offset of the Gabor, a mask consistent of 6 concentric circles (line color: black; radii = 1.25, 2.00, 2.75, 3.50, 4.25, 5.00) was presented. This mask had a duration of 300 ms and was presented in order to eliminate the possibility of subjects experiencing an afterimage. Following a blank period of 400ms, subjects were presented with a white Gabor patch (3 cycles/degree, only one strip visible, random initial orientation) that they rotated via button press (up and down arrow, resolution = 1 degree) until they considered the orientation of the white Gabor patch indicator to match that of the target Gabor. Subjects logged a response via button press. The inter-trial interval was set to 1 second, and participants completed 200 trials per block. The experiment consisted of 3 blocks; the first was without feedback, as described above. In the second and third blocks participants were given feedback by superimposing the target Gabor and the orientation reported during the inter-trial interval. Participants were given approximately 5 minutes rest between blocks. All stimuli were generated and rendered using C++ Open Graphics Library (OpenGL).

### Data Analyses

We model subjects as a generic estimator *T* of the orientation parameter *θ*, and thus the responses as independent samples of *T*. The bias and variance of the estimator *T* is a function of *θ* defined by *b*(*θ*) = *Eθ*[*T*] − *θ*, and *σ*^2^(*θ*) = *σ*^2^*θ* [*T*], respectively. The lower bound of Fisher information (FI) as a function of l is given by Cramer-Rao Lower Bound (**Fig. 3**):

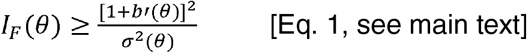

For our analysis, we assumed a tight bound (e.g., both Bayesian and maximum likelihood estimator can attain the bound asymptotically, also see Fig. 3), which allows us to extract FI directly from data (**Fig. 3, 4C**). Furthermore, with an efficient coding assumption that the square-root of FI is directly proportional to the prior subjects adopt:

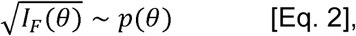

we can thus estimate the prior distribution by calculating the normalized square-root of FI:

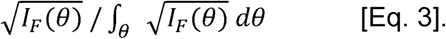

To quantify the changes in encoding of subjects in the experiment, we parameterized the square-root of FI wi λ and □:

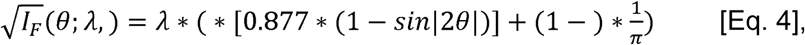

where *λ* determines the amount of total (square-root of) FI, and controls the shape, or “allocation” of FI, effectively the prior distribution, by mixing a natural orientation prior distribution with a uniform distribution (**Fig. 4C**). Note the constants are such that the prior part of the equation normalizes properly to 1.

To recover λ and □, we calculate the predicted bias:

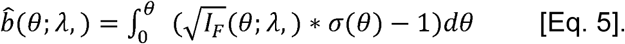

We find 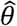 and ^ that give rise to the best fit to the observed bias *b*(*θ*)(**Fig. S1**):

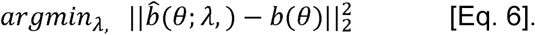

Note that here we choose to fit the bias pattern *b*(*θ*), mainly to avoid the noisy derivative, *b′*(*θ*) in the Cramér-Rao Lower Bound (and introduce an integration step instead).

**Fig. 4D, E** shows the parameters estimated for the combined subject, while the regression analysis presented in **Fig. 5** is based on parameters estimated with the same procedure applied to each individual subject. All tests and p-values (except for the regression in **Fig. 5**) reported are based on the distribution (intervals) of the sample statistics (e.g., mean, variance, model parameters *ω* and *λ*) across 5,000 bootstrap runs (Efron & Tibshirani, 1994). Tests are one-sided unless specified otherwise in the main text.

## Acknowledgements

We thank J. Patterson and A. Rosenberg for piloting some of the stimuli used in the present experiments and H. Park for participation in data collection. This work was supported by the Simons Foundation, SFARI Grant 396921.

## Author contributions

D.E.A. designed and supervised the experiment. J.P.N. and L.Q.Z. analyzed data. L.Q.Z. and A.A.S. performed the modeling. J.P.N., L.Q.Z., A.A.S. and D.E.A. wrote manuscript.

## Supplementary Figures

**Fig. S1:**
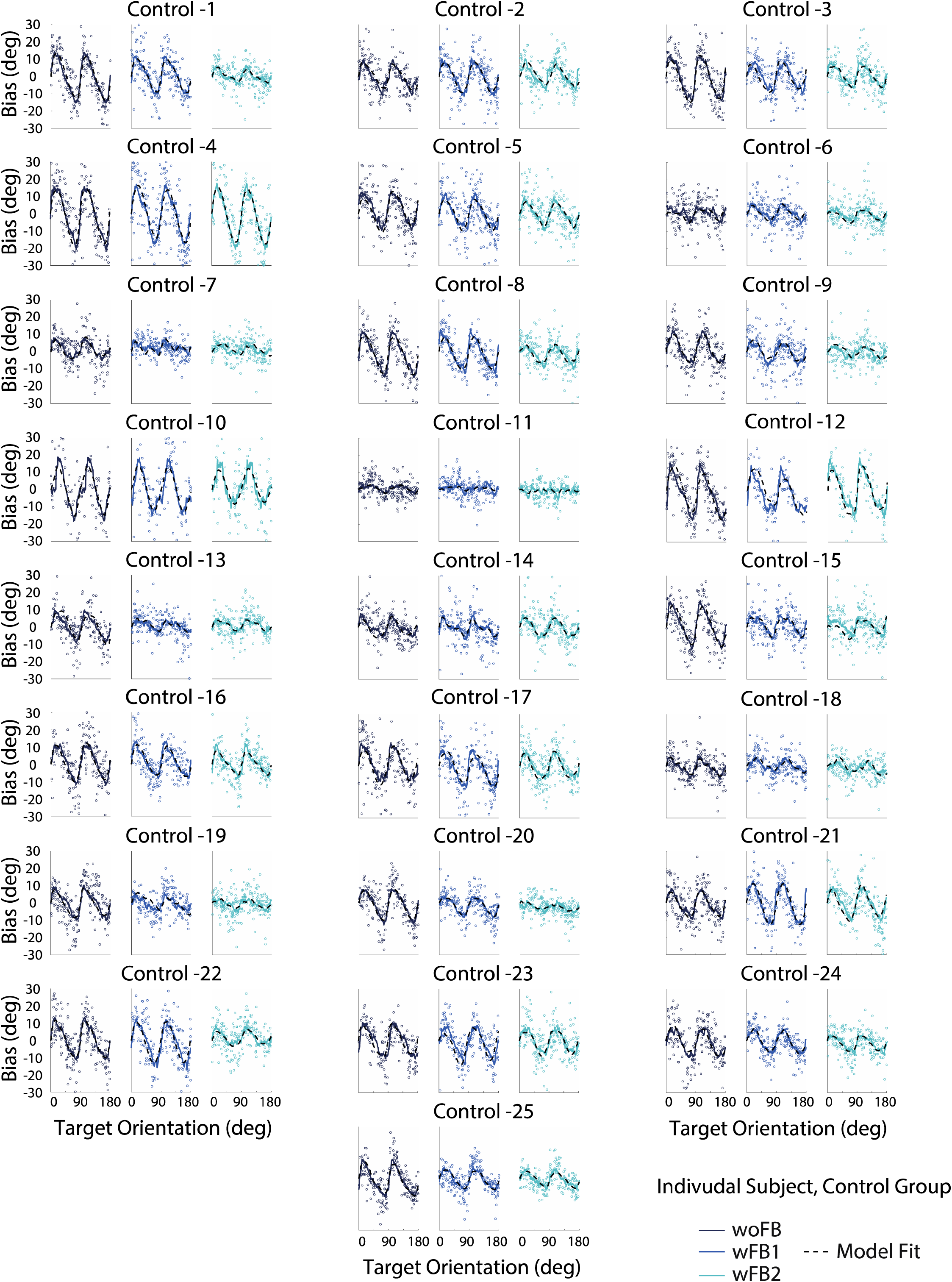
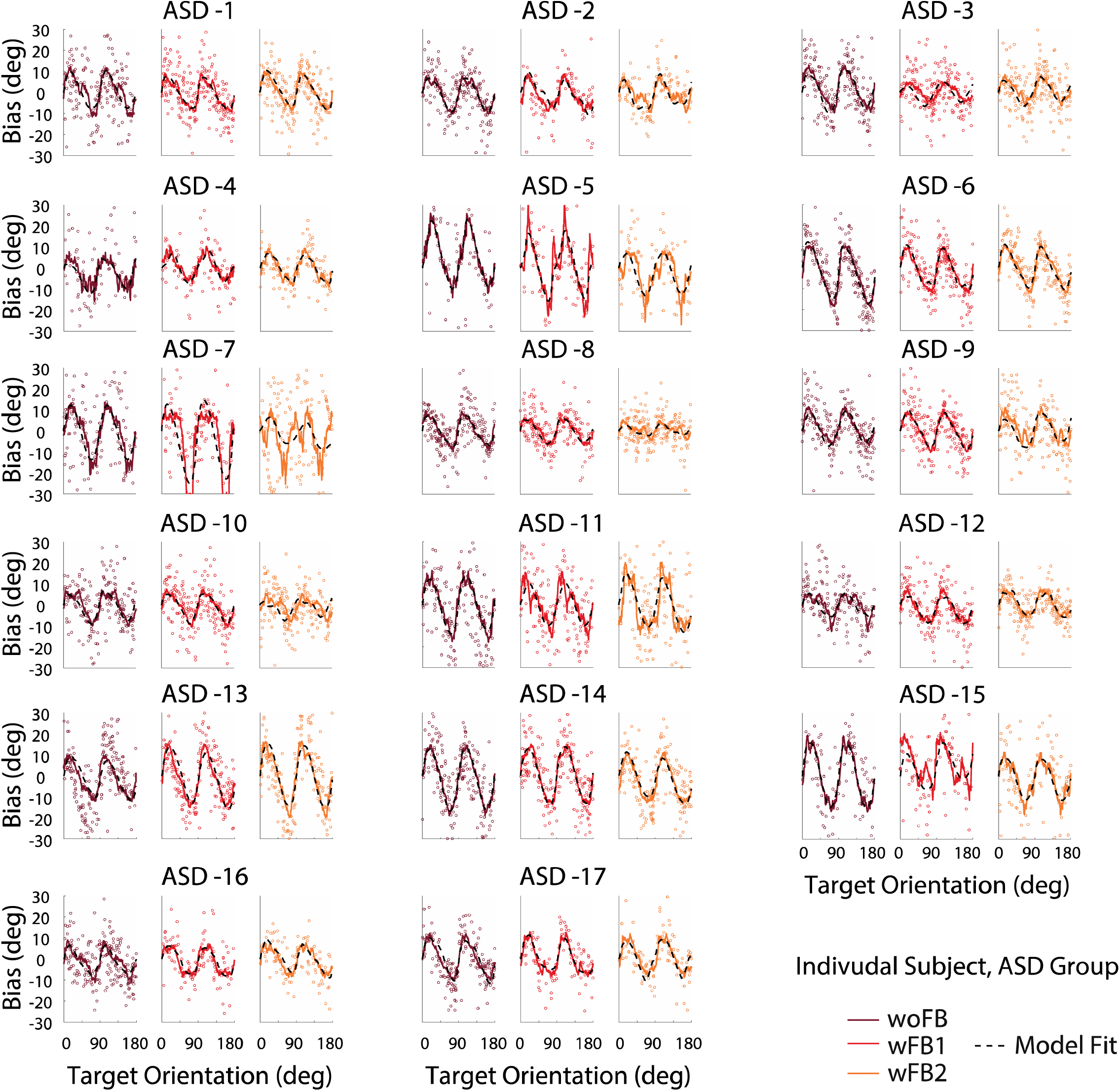
Scatter plot of the response data for individual subject, and the model fits to the average bias pattern. Note that we enforced the bias pattern to have a period of 90° to increase the reliability of the fits.

**Fig. S2:**
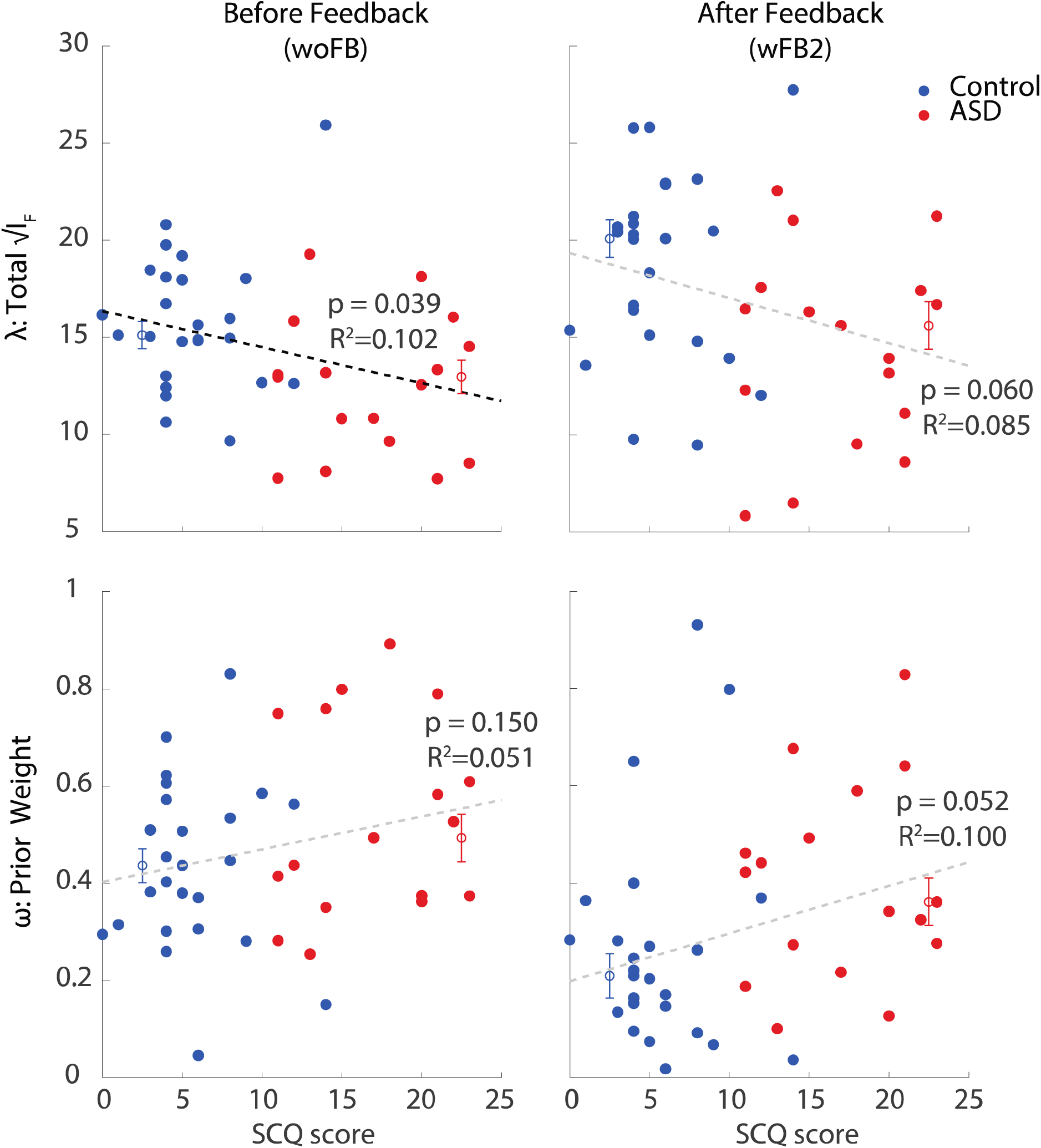
Correlation of extracted parameter with SCQ scores.

**Fig. S3:**
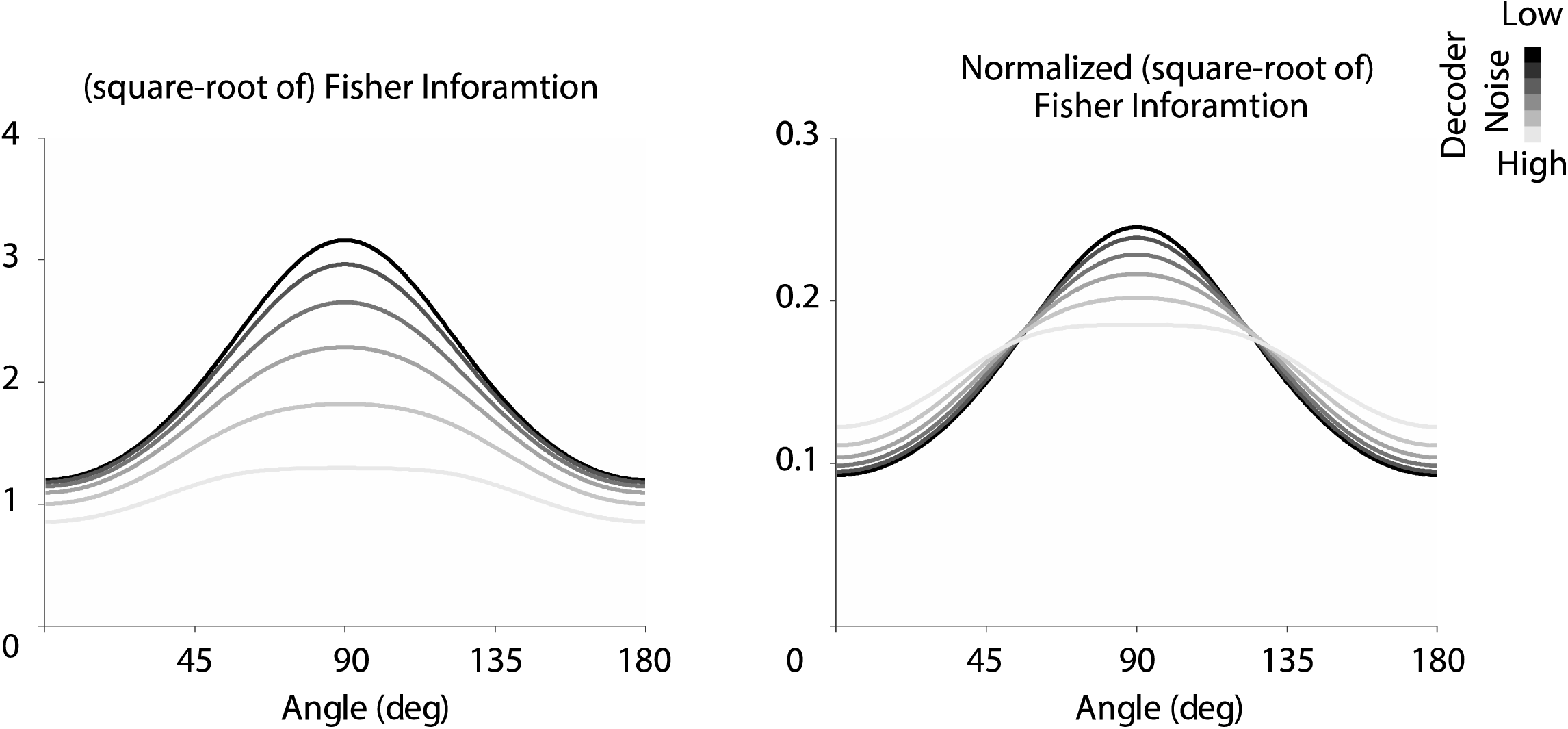
Extracted pattern of FI using Cramer-Rao lower bound when there is a stimulus-independent inefficiency (homogeneous noise in this case) added to the decoder.

